# Mitochondrial control of microglial phagocytosis in Alzheimer’s disease

**DOI:** 10.1101/2021.12.01.469111

**Authors:** Lauren H. Fairley, Kei Onn Lai, Jia Hui Wong, Anselm Vincent Salvatore, Giuseppe D’Agostino, Xiaoting Wu, Anusha Jayaraman, Sarah R. Langley, Christiane Ruedl, Anna M. Barron

## Abstract

Microglial phagocytosis is an energetically demanding process that plays a critical role in the removal of toxic aggregates of beta amyloid (Aβ) in Alzheimer’s disease (AD). Recent evidence indicates that metabolic programming may breakdown in microglia in AD, thereby disrupting this important protective function. The mechanisms coordinating mitochondrial metabolism to fuel phagocytosis in microglia remain poorly understood, however. Here we demonstrate that mitochondrial displacement of the glucose metabolizing enzyme, hexokinase-II (HK) regulates microglial metabolism and phagocytosis, and that deletion of the translocator protein (TSPO) inhibits this. TSPO is a PET-visible inflammatory biomarker and therapeutic target in AD, previously shown to regulate microglial metabolism via an unknown mechanism. Using RNAseq and proteomic analyses, we found TSPO function in the brain to be linked with the regulation of mitochondrial bioenergetics, lipid metabolism and phagocytosis. In cultured microglia, TSPO deletion was associated with elevated mitochondrial recruitment of HK, which was associated with a switch to non-oxidative glucose metabolism, reduced mitochondrial energy production, lipid storage and impaired phagocytosis. Consistent with *in vitro* findings, TSPO expression was also associated with phagocytic microglia in both AD brain and AD mice. Conversely, TSPO deletion in AD mice reduced phagocytic microglia and exacerbated amyloid accumulation. Based on these findings we propose that microglial TSPO functions as an immunometabolic brake via regulation of mitochondrial HK recruitment, preventing hyperglycolysis and promoting phagocytosis in AD. Further, we demonstrate that targeting mitochondrial HK may offer a novel immunotherapeutic approach to promote microglial phagocytosis in AD.

## Introduction

Mitochondria are emerging as the command centers of innate immune responses, controlling inflammatory responses via metabolic programming. This programming is known as immuno-metabolism (reviewed 1). Recent evidence indicates that in Alzheimer’s disease (AD), metabolic programming breaks down in microglia, which are the resident innate immune cells of the brain, thereby disrupting important protective functions of these cells such as phagocytosis (2–5). Microglial phagocytosis plays a key role in the clearance of toxic beta amyloid (Aβ), aggregations of which are a hallmark of AD (6). The mechanisms coordinating mitochondrial metabolism to fuel phagocytosis, which are disrupted in AD, remain poorly understood. Here we investigate the role of the mitochondrial translocator protein (TSPO) complex in microglial metabolic programming and phagocytosis in AD-related inflammation.

TSPO is a PET-visible inflammatory biomarker (7–9) and therapeutic target in AD (10–14). Microglial TSPO expression is upregulated in AD, correlating with the distribution of pathological aggregates of Aβ and tauopathy (9, 15, 16). However, despite being widely regarded as a marker of microglial activation, the specific immunophenotype associated with TSPO, and whether TSPO modulates beneficial or detrimental microglial functions in AD has not been determined. Further, the function of TSPO in microglial immune responses and AD pathogenesis remains unclear. In mouse models of AD, we and others have shown that TSPO-targeting drugs reduce Aβ accumulation and neuronal death (10, 14, 17). While TSPO has been implicated in a range of microglial inflammatory responses, including cytokine secretion, proliferation, motility and phagocytosis (18–20), the molecular mechanism underlying the protective effects of TSPO-targeted drugs in AD are not understood.

One possibility is that TSPO regulates microglial inflammatory responses via metabolic reprogramming, although how this might occur is unknown (21–25). Pro-inflammatory activation of microglia is dependent upon a switch from energy production via mitochondrial oxidative phosphorylation (OXPHOS) to non-oxidative glucose metabolism via glycolysis (4, 5, 26–28). Increased glycolysis, or hyperglycolysis, is associated with aging and AD, linked to compromised phagocytosis and Aβ engulfment (reviewed in 1, 29-35). This switch to aerobic glycolysis is known as the Warburg effect, originally described in cancer. In cancer, the balance of oxidative and non-oxidative glucose metabolism is coordinated by the voltage-dependent anion channel (VDAC), which forms part of a mitochondrial complex with TSPO (36–38).

VDAC is a mitochondrial hub protein and the major channel for supply of mitochondrial substrates to fuel OXPHOS. In other cell types, binding of the VDAC-complex to the glycolytic enzyme, hexokinase II (HK), allosterically increases its activity to promote glycolysis whilst inhibiting OXPHOS (39). Although currently there is little agreement on what interactors form the functional complex with TSPO in the neuroimmunological context, the same HK isoform is expressed by microglia and is known to play a critical role in mediating inflammatory activation (40).

Here, to determine the TSPO-associated immunological signature in a model of neuroinflammation, we examined the effect of TSPO deletion (TSPO-KO) on microglial function in models of AD-related inflammation. These findings provide new evidence that microglial metabolic programming and phagocytic function is regulated via changes in subcellular localization of the key metabolic enzyme, HK. Based on these findings we propose that TSPO functions as an immunometabolic brake via regulation of mitochondrial HK recruitment, preventing hyperglycolysis and promoting phagocytosis in AD.

## Methods

### Animals and treatments

C57BL/6 (wild type, WT), homozygous TSPO-KO (41), APP NL-G-F knock-in (APP-KI) (42) and APP-KI x TSPO-KO mice were bred and maintained under specific pathogen-free conditions at Lee Kong Chian School of Medicine Animal Research Facilities with food and water available *ad libitum*. All experiments were carried out in accordance with the National Advisory Committee for Laboratory Animal Research guidelines and approved by the by the NTU Institutional Animal Care and Use Committee (IACUC# A0384).

For RNAseq and microglial lipid droplet experiments, 3 month-old WT (C57BL/6J) and TSPO-KO mice were injected with phosphate buffered saline (PBS, control) or LPS (Lipopolysaccharide from *E. coli*, Sigma Aldrich; 1 mg/kg body weight i.p.) daily for four consecutive days.

For tissue collection in all experiments, mice were deeply anesthetized with sodium pentobarbital (200 mg/kg i.p.) and transcardially perfused with 0.9% saline. Brains were rapidly removed and either snap frozen on 2-methylbutane on dry ice and stored at −80°C for RNA extraction, submersion fixed overnight in 4% paraformaldehyde (PFA) in PBS for immunohistochemistry (IHC), or placed in IMDM (2% FBS) for cell dissociation and flow cytometry analysis. Fixed tissue was cryoprotected in 30% sucrose prior to freezing and sectioning. Sections were cut at 20 μm by cryostat, then stored in cryoprotectant solution at −30°C until use.

### RNAseq data generation & analysis

Total RNA was extracted from WT and TSPO-KO mouse hippocampus (see supplementary methods) and OligodT mRNAseq stranded library preparation were conducted by Novogene AIT using NEBNext Ultra library preparation kit and sequenced using Illumina HiSeqTM. Quality control of the FASTQ files were performed using FASTQC in Ubuntu 18.04 environment. Raw sequence were aligned using pseudo aligner Salmon according to Mus musculus genome Ensembl release 99. Reads of individual transcripts were loaded onto Rstudio and differential expression analysis were performed using DESEQ2 and significance tested using FDR Bonferonni correction for multiple testing (43). Log_2_-fold change (LFC) values were shrunk using ashr package and thresholded at LFC ≤ 0.5 or LFC ≥ 0.5 (44). Volcano plots of RNAseq were constructed using Enhanced Volcano package (45) in an Rstudio environment and heatmap was plotted using Complex Heatmap package (46). For gene set enrichment, fGSEA package (47) was used and the Wald statistic column output (stat) of the differentially expressed genes were run against MsigDB collection’s C5 (Gene Ontology) Geneset of Musmusculus (number of permutations of independence testing = 10000). fGSEA results were pruned using an in-house script to remove repetitive terms and prevent over-correction of nominal p-values during multiple test correction (48). FDR was calculated for each Gene ontology subcategory to correct for multiple testing (either belonging to Biological Process, Molecular Function or Cellular Function). Significantly enriched terms were curated by the experimenter according to functional clusters then ranked within functional clusters by FDR and NES. The full analysis pipelines are available at this link https://github.com/Neurobiology-Aging-and-Disease-Lab/Fairley-Lai-et-al-.

### Immunoprecipitation mass-spectrometry (IP-MS)

Proteins were IP from mouse brain homogenate using rabbit anti-TSPO antibody (Abcam 109497; see supplementary methods for details) and LC-MS and peptide detection was provided by Proteomics and Mass Spectrometry Facility in School of Biological Sciences, Nanyang Technological University. Empai scores of identified proteins were imported to R environment version 4.1 and converted to log2 format using DEP bioconductor R package (49) Version 3.13. For identification of candidate TSPO interactors, WT and APP-KI data was analysed together as “positive samples”, and TSPOKO-/- data is taken as “background”.

Only proteins detected in at least 5 of the 6 positive samples were considered for analysis. Missing at Random (MAR) values for “positive samples” were imputed using KNN and Missing Not At Random (MNAR) values in “background” samples are imputed using Minimum Probability method (50). Imputation was performed in R environment using imputeLCMD (51) and impute package (52). Candidate interactors were defined as significantly enriched in positive samples relative background (FDR < 10%, fold change > 2) determined by differential enrichment analysis using Bioconductor R package Limma version 3.13 (53). Network and functional enrichment analysis of identified candidate interactors was carried out using Cytoscape (54), InTact (55) and StringApp plugins version 3.8 (56). The STRING network was mapped using a confidence score of 0.4 under “Mus musculus” and the functional enrichment retrieved using KEGG and Reactome pathways databases.

### Mitochondrial bioenergetics in primary microglia

Mitochondrial bioenergetics in cultured primary microglia were measured using the XF Cell Mito Stress Test Kit (#103015-100, Seahorse Biosciences, North Billerica, MA) according to methods described in the XFe96 Extracellular Flux Analyzer User Manual. For detailed description see supplementary methods.

### TMRE

Primary microglia were labelled with TMRE (200nM, Abcam ab113852) and fluorescent intensity measured by microplate reader (excitation: 549nm, emission: 575nm). Data was normalized to total cell number measured using DAPI.

### Hexokinase activity assay

Hexokinase activity was measured in primary microglial culture cell lysates using Hexokinase Activity Assay Kit (Colorimetric) (Abcam, ab136957) according to the manufacturer’s instructions Data was normalized to total protein concentration.

### Intracellular ATP live cell imaging

To measure intracellular ATP levels, WT and TSPO-KO primary microglia, cells were stained with MitoTracker™ Deep Red FM (Cat. No.: M22426, ThermoFisher Scientific) and ATP-Red 1 (Cat. No.: HY-U00451, MedChem Express) and imaged by confocal microscopy.

### *In Vitro* Phagocytosis Assay

Cultured BV-2 cells or primary microglia were incubated with Latex beads (Sigma Aldrich, L4530; ∼100 beads per cell), or 500 nM fluorescently-labelled oligomeric Aβ (o-Aβ) for 2 hrs. To displace mitochondrial HK, Hexokinase II VDAC Binding Domain Peptide (Hkp, 2.5uM; Merck, 376816) or vehicle (water solvent) was added simultaneously with bead. Bead or o-Aβ uptake was measured by flow cytometry or analysed in fixed cells by microscopy. Phagocytic efficiency was measured by counting number of total cells with at least one internalized bead. For experiments in which LPS was used, cells were stimulated with LPS 100 ng/mL for 2 hrs prior to incubation with phagocytic stimuli.

### Immunocytochemistry

Cells were labelled with polyclonal rabbit anti-hexokinase (Abcam, ab227198), and monoclonal mouse anti-ATPB (Abcam, ab14730), followed by fluorescent-conjugated secondary antibodies and counterstained with Hoechst 33342 (ThermoFisher 62249, 1 μg/mL in PBS). F-actin was labelled prior to cell fixation using Phalloidin-647 (Abcam ab176759). Images were captured using a confocal microscope (Zeiss Laser Scanning Microscope-780 upright microscope) and analysed using Imaris 9.2.0.

### Flow Cytometry

Isolation of myeloid cell populations was performed as previously described (57). Isolated cells were incubated with anti-mouse CD16/32 in PBS for 15 min to block the Fc receptor. Cells were then washed with PBS and resuspended in the desired antibody mix diluted in PBS and stained for 30 min at 4°C protected from light. Antibody details are provided in supplementary table 1. For staining with intracellular markers, cells were fixed in 4% PFA and permeabilized with 0.1% Triton X-100 for 5 min prior to staining. For lipid droplet analysis, cells were not permeablized and were incubated with BODIPY –FLC12 (1:1000) for 10 min at RT. Cell viability was assessed using propidium iodide. Single-cell suspensions were then analyzed using a LSRII flow cytometer (Becton Dickinson) and data were analyzed with FlowJo software (TreeStar).

### Immunohistochemistry

Mouse brain tissue sections were permeabilized in 0.1% Triton X-100 for 5 mins. Sections were blocked in 5% BSA + 0.2% Triton-X-100 for 1.5 hours to prevent non-specific binding, followed by overnight immunolabeling at 4°C with primary antibodies. After incubation with the primary antibody, sections were washed three times in PBS for 5 min and then incubated with secondary antibodies for 1.5 hr at room temperature. After washing with PBS and water, they were mounted on microscope slides using Fluroshield with DAPI (Sigma F6057) and stored at 4°C for imaging.

For immunofluorescence using human post-mortem tissue (see supplementary methods and suppl. Table 2 for sample details), 10 µm thick snap-frozen tissue sections were fixed with 100% methanol at −20°C, blocked with 2.5% goat serum, and incubated with primary antibodies overnight. The sections were incubated with the appropriate secondary antibody conjugated with a fluorochrome and mounted with Vectashield® Vibrance^TM^ Antifade Mounting medium with DAPI (Vector Laboratories) for nuclei counterstaining. The primary antibodies used were rabbit polyclonal anti-GFAP (Abcam, ab7260), rabbit polyclonal anti-TMEM199 C-terminal (Abcam, ab185333), anti-Iba1 (WAKO, 019-19741), anti-PBR (Abcam, ab199779), mouse monoclonal anti-Aβ (Merck, MABN639).

Images were captured using a confocal microscope (Zeiss Laser Scanning Microscope-780 upright microscope) with Zeiss ZEN imaging software. Image analyses were performed in a user-blinded manner using ImageJ and de-noised using background subtraction rolling ball radius of 50 pixels. Separate high-resolution z-stacks imaged at 63x resolution, with an optimal step size, were added to determine the levels of TSPO in Iba1-positive microglia proximal to regions of plaque contact (proximal; within 10 μm of a plaque) compared to regions where there was no plaque contact (distal; within 80 μm of a plaque). Mean TSPO expression was divided by the %Area of IBA-1 signal to account for differences in microglia numbers in regions proximal versus distal to plaques.

### Statistical Analysis

Statistical Package for Social Sciences (SPSS Inc), GraphPad Prism 8.2.1 (GraphPad Software) or the online server at http://www.estimationstats.com (58) was used for statistical analyses and generation of plots. For comparison of two groups, a two-sided permutation t test was used and effect sizes were estimated by calculating bootstrapped 95% confidence intervals (CI) and are displayed with the bootstrap distribution of the mean (58). Levinès and Shapiro-Wilk tests were used to assess homoscedasticity and normality. For comparisons of more than two groups, One- or Two-Way ANOVA was used. Significant factors or interactions were further explored using Bonferroni post-hoc comparisons.

## Results

### TSPO deletion alters transcriptional profiles in phagocytosis, mitochondrial metabolism and lipid biosynthesis pathways in neuroinflammation

To determine the effect of TSPO deletion on neuroinflammatory responses, LPS-induced transcriptional profiles in the hippocampus of WT and TSPO-KO mice were compared by RNA-seq. Under control conditions, little effect of TSPO-KO was observed on transcription, with only 27 significantly differentially expressed genes (DEGs, FDR < 0.05, shrunkenLFC > shrunken LFC ≥0.5 or ≤-0.5; Fig. 1A). In contrast, following stimulation with LPS, robust transcriptional differences were observed between TSPO-KO and WT mice, with 795 significantly DEGs, supporting the notion that TSPO plays a role in inflammatory responses in the brain (545 downregulated, 250 upregulated; Fig. 1A & Suppl. Table 3). Functional gene set enrichment analysis (FGSEA) comparing LPS-stimulated TSPO-KO and WT mice showed downregulation of immune and inflammatory pathways, and upregulation of functional clusters of mitochondrial metabolism and lipid metabolism pathways (Fig. 1B & Supp. Table 4).

**Fig. 1.**
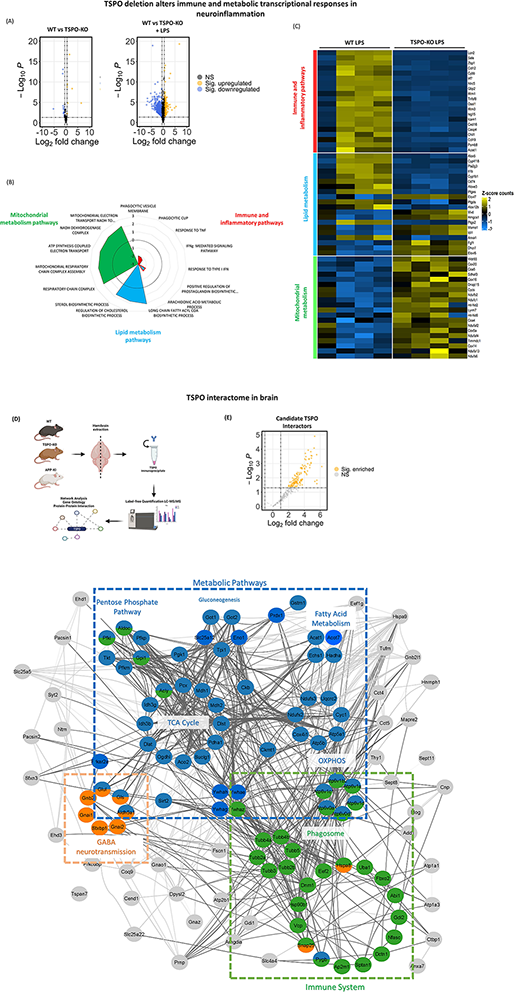
TSPO function associated with mitochondrial bioenergetics, lipid metabolism and phagocytosis pathways in AD-related neuroinflammation. **(A)** Volcano plot showing differentially expressed genes in hippocampus of WT versus TSPO-KO mice (n = 4/group). (FDR < 0.05; NS, non-significant). **(B)** Normalized enrichment score (NES) of top enriched GO pathways related to immune and inflammatory responses, mitochondrial and lipid metabolism identified by FGSEA (FDR < 0.05). **(C)** Heat map of mean gene expression for selected immune-, mitochondrial and lipid metabolism in WT and TSPO-KO mice. **(D)** Schematic of proteomic TSPO interactome analysis in brain. TSPO complexes were isolated via immunoprecipitation from wildtype and APP-KI mouse brains, using TSPO-KO as a background control. Enriched proteins were identified by LC-MS/MS and the interactome network mapped using bioinformatics approaches. Created with BioRender.com. **(E)** Volcano plot showing proteins identified by TSPO IP-MS, with candidate TSPO interactors (orange data points) identified as greater than 2-fold significantly enriched compared to background (TSPO-KO) (FDR < 0.1). **(F)** Brain TSPO interactome protein-protein interaction network showing immune system and metabolic pathways functional clusters (Functional enrichment FDR < 0.05). Abbreviations:

Top downregulated immune-related processes were phagocytosis, and responses to tumor necrosis factor (TNF) and interferon (IFN) signaling. Downregulation of phagocytosis-related pathways included genes involved in phagocytic cup formation and closure (INPP5B, IRGM1, DNM2), phagosome acidification and respiratory burst (SLC11A1, ATP6V0E, CYB, NCF1,2,4), phagosolysosome assembly and maturation (CORO1A, RAB, RAC, VAV), and expression of phagocytosis receptors (FcγR -FCGR, Bin2, CR1L, CR3, ITGAM, ITGB2, TLR1, TLR2, CD14, MSR1, MARCO) (Figure 1B, 1C). Key genes in downregulated TNF and IFN signaling pathways in TSPO-KO mice included TNF, TNFSF-10 and TNFRSF (13b,14). Genes involved in antigen processing and presentation (TAP, H2-M3, H2-T23) and complement cascade (C3, C5, Cf b) were also downregulated. These findings support a key role for TSPO in neuroinflammatory responses, particularly phagocytosis and cytokine signaling.

Supporting existing consensus on TSPO’s role as a mitochondrial metabolic target, upregulation of genes encoding subunits of mitochondrial complexes I, II and IV was observed (ND, NDUFA-C, NDUFS, SDH, COX). Likewise, cholesterol and fatty acid biosynthesis genes were upregulated (ACSL, ACSBG1, CH25H, FASN, ELOVL, MVD), while synthesis and metabolism of the inflammatory eicanosinoids was downregulated (ALOXE3, ALOX5, ALOX5AP, PTGES) in LPS stimulated TSPOKO mice (Fig. 1B, C). Clusters of other functionally enriched pathways included downregulation of transcription and translational responses, apoptosis and cell-death related pathways, and upregulation of terms related to nervous system development, myelination, ion homeostasis and excitatory/inhibitory neurotransmission, including GABAergic neurotransmission related pathways.

### Proteomic TSPO interactome network in AD

To gain insight into the protein-protein interactions and molecular pathways underlying TSPO-complex function in inflammatory responses in the brain, the TSPO interactome was captured by co-IP-MS under control and AD inflammatory conditions using WT and APP-KI mouse brain. A total of 124 proteins were identified as candidate TSPO interactors, significantly enriched relative to TSPO-KO background (Suppl. Table 5).

Analysis of the candidate TSPO-interactors for known physical protein-protein interactions using IntAct database revealed the 14-3-3 family scaffold adaptor and chaperone proteins, Ywhae, Ywhab and Ywhaz, to interact with the majority the identified TSPO-interactors (87.5 %, 63.3 % % and 48.3 % of TSPO-interactors, respectively; FDR < 0.0001). The 14-3-3 adaptor proteins are metabolic regulators (59) and have been previously shown to interact with the TSPO-complex in leydig cells (60, 61). Other previously reported TSPO interactors identified in the brain TSPO interactome included ATP synthase (Atp5a1), mitochondrial solute carrier family 25 (Slc25a5) and mitochondrial creatine kinase (Ckmt1).

The brain TSPO interactome network was mapped using STRING network analysis (Fig. 1D-F). Of the 124 identified candidate interactors, 44 were enriched from the mitochondrial compartment. Functional enrichment of metabolism pathways and immune system and was observed in the TSPO interactome network (FDR < 0.05; Fig. 1F, Suppl. Table 6). Metabolism pathways included functional clusters in pentose phosphate pathway, gluconeogenesis/glycolysis, TCA cycle, oxidative phosphorylation and fatty acid metabolism. Immune system functional clusters included phagosome-related networks, including vacuolar ATPase involved in phagosome acidification and cytoskeletal elements involved in phagosome transport. Functional enrichment of terms relating to neurotransmission (including GABA synthesis, release, reuptake and degradation) and neurodegenerative disease (including Alzheimer disease, Parkinson disease, Prion disease, Huntington disease, Amylotrophic lateral sclerosis) were also observed.

### TSPO-KO modulates mitochondrial hexokinase binding and induces a switch to non-oxidative cellular bioenergetics

Since our proteomic data implicated TSPO in metabolism pathways and immune function, we examined the effect of TSPO-KO on cellular bioenergetics in cultured microglia. TSPO-KO microglia exhibited impaired mitochondrial OXPHOS and reduced mitochondrial ATP levels (Fig. 2A - C). A marked reduction in basal respiration (t-test: CI = −77.6, −60.3; *p* < 0.0001), maximal respiration (t-test: CI = −1.05e+02, −48.2; *p* = 0.0014) and ATP production (t-test: CI = −53.0, −42.2; .*p* < 0.0001) as by measured oxygen consumption rate (OCR) was observed in TSPO-KO microglial cultures (Fig. 2B). Reduced mitochondrial, but increased cytosolic ATP signals in TSPO-KO microglia was confirmed using the fluorescent ATP probe, ATP-red (Fig. 2C). Reduced mitochondrial membrane potential was also observed in TSPO-KO microglia (CI = −0.795, −0.508, *p* = 0.0034; n = 4 – 6 wells), consistent with respiratory chain defects and reduced OXPHOS (Suppl. Fig. 1A). Metabolic impairment in TSPO-KO microglia was coupled with elevated activity of the rate-limiting glucose metabolizing enzyme, hexokinase-II (HK) (HK activity t-test: CI = 0.63, 5.76, *p* = 0.0252; n = 9-10 wells/group, average of two independent experiments; Fig. 2D). HK is a known interactor of the TSPO-VDAC complex, exhibiting increased enzymatic activity and inducing a switch from OXPHOS to non-oxidative glucose metabolism in the cytosol upon binding to mitochondrial VDAC (62). Consistent with this, increased mitochondrial HK binding was observed in TSPO-KO microglia (t-test: CI = 6.97 - 18.5, *p* = 0.0002; n = 40 individual cells/group, average of two independent experiments; Fig 2E, FI). Interestingly, this increase in mitochondrial HK binding was observed in TSPO-KO microglia despite a modest but significant decrease in HK mRNA expression in mouse brain following LPS stimulation (LFC: −0.548, *p* = 0.013). A modest reduction in mitochondrial content as measured by footprint and increase in mitochondrial fragmentation was also observed in TSPO-KO microglia under baseline conditions (mitochondrial footprint *t-*test: CI = −7.47, −1.9; *p* = 0.001; fragmentation t-test: CI = 0.0144 - 0.12, *p* = 0.0196; Suppl. Fig. 1B, C).

**Fig. 2.**
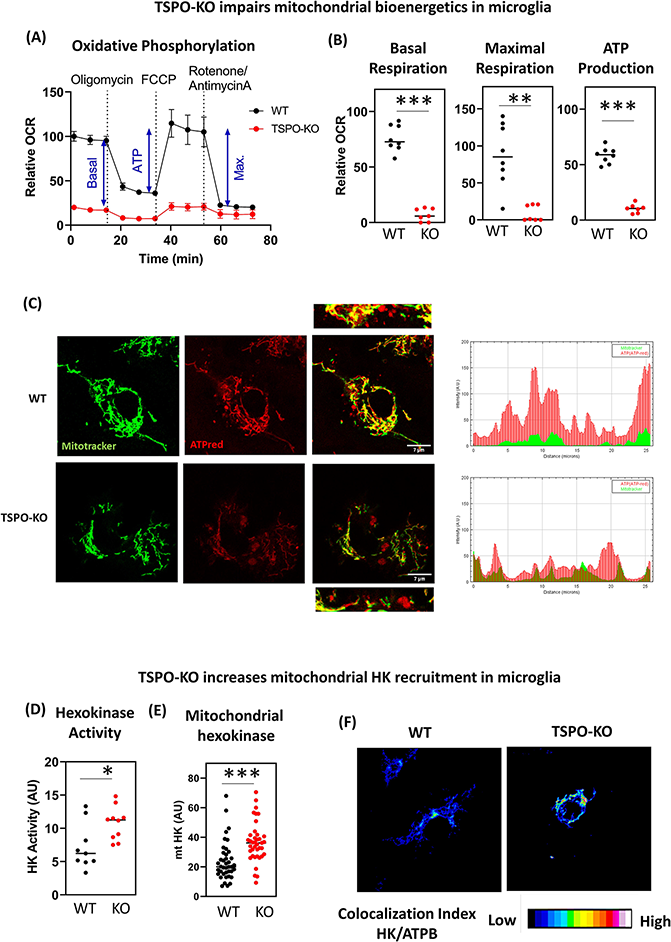
TSPO deletion induces mitochondrial hexokinase binding and impairs mitochondrial bioenergetics in microglia. (**A**) Mitostress test of WT and TSPO-KO cultured microglia showing mean ± SEM oxygen consumption rate (OCR) normalized to WT. Basal respiration (basal), ATP production (ATP) and maximal respiration (max.) indicated on WT curve. n = 7-8 wells/group, average of two independent experiments. (**B**) Quantification of basal respiration, maximal respiration and ATP production calculated from Mitostress assay. Dots represent individual wells. (**C**) Representative image of mitotracker (green) and intracellular ATP signals (red) in cultured WT and TSPO-KO primary microglia. ATP colocalization with mitochondria can be observed in the merged panel (yellow). Kymograph indicates ATPred and mitotracker intensity in line indicated on merged image. (**D**) Hexokinase activity measured in cell lysates. Dots represent individual wells (n = 9 – 10/group). (**E**) Mitochondrial hexokinase binding (mtHK) measured by median hexokinase immunoreactive intensity within mitochondrial volumes determined using ATPB staining in WT and TSPO-KO microglial cultures. Dots represent individual cells (n = 40/group). (**F**) Representative confocal images of mitochondrial hexokinase binding shown as colocalization index of hexokinase relative to mitochondrial marker, ATPB. AU: arbitrary units; OCR: oxygen consumption rate. \**p* < 0.05. *\*\* p* < 0.002. *\*\*\*p* < 0.0001.

### TSPO deletion induces microglial lipid droplet accumulation

Since with metabolic reprogramming from OXPHOS to glycolysis is associated with lipid droplet formation in aging and disease (63, 64), we investigated lipid droplet accumulation in primary microglial cultures using the neutral lipid marker, BODIPY –FLC12 (BD). Consistent with the transcriptomic and proteomic data suggesting a role for TSPO in fatty acid synthesis and metabolism, primary microglia isolated from TSPO-KO mice exhibited enhanced lipid-droplet accumulation compared to WT primary microglia (Two way ANOVA; Main Effect (Genotype): F (_1,30_) = 45.668; *p <* 0.001), and LPS markedly exacerbated lipid droplet accumulation in both cultured TSPO-KO and WT microglia (Two way ANOVA; Main Effect (Treatment): F _(1,30)_ = 23.018; *p <* 0.001; n = 9/group, average of 3 independent experiments, Fig. 3A, B). In contrast, TSPO ligand Ro5-4864 attenuated LPS-induced lipid droplet accumulation in immortalized microglial BV2 cells (Two way ANOVA; Interaction Effect (Treatment x LPS): F _(1,32)_ = 52.668; *p <* 0.001; Pairwise Comparison (Ro5 + LPS v Vehicle + LPS): *p <* 0.001; n = 9/group, average of 3 independent experiments; Fig. 3C). In mouse brain, significantly lower TSPO expression was observed in the LPS-induced lipid droplet accumulating microglial subpopulation (t-test: *CI* = −7.78, - 2.19, *p* < 0.001; n = 3 mice; Suppl. Fig. 1E). A recent study has associated lipid droplet accumulation with phagocytosis-impaired, dysfunctional microglial phenotypes that develop in aging (65). Consistent with this, we observed reduced levels of the lipid droplet marker, perilipin 2 (PLIN2), in phagocytic compared to non-phagocytic microglia (*t*-test: CI= - 7.07e+03, −2.93e+03; *p* < 0.0001; n = 3/group; Suppl. Fig 1F).

**Fig. 3.**
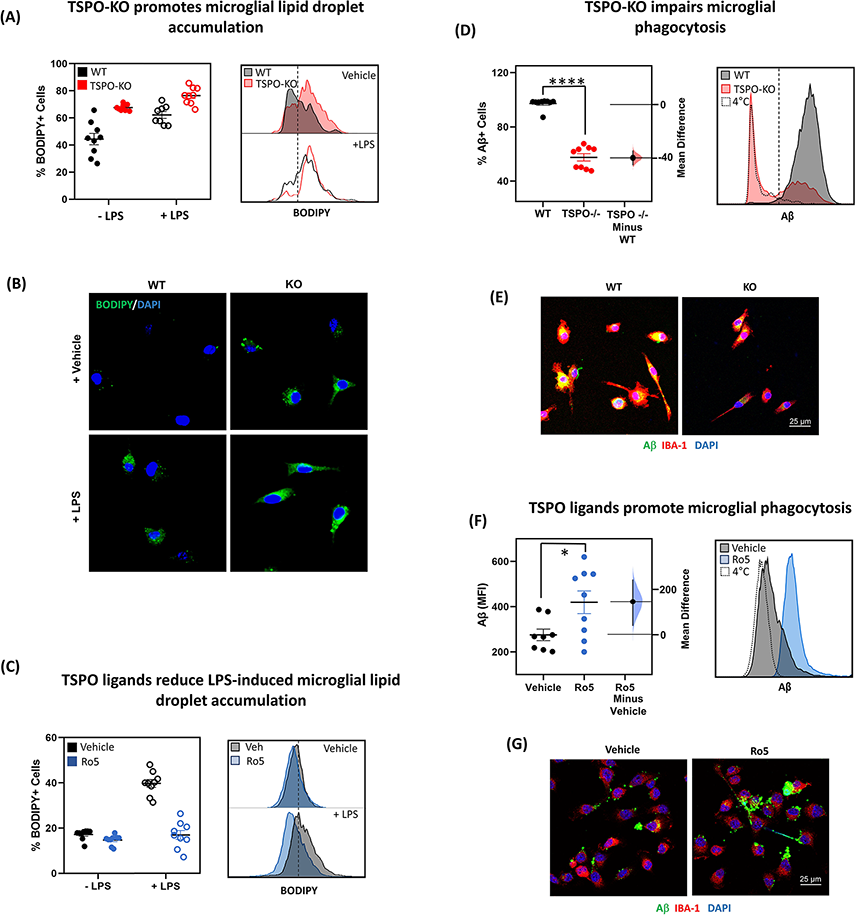
TSPO deletion promotes lipid droplet accumulation and impairs phagocytosis in microglia. (**A**) TSPO deletion promotes microglial lipid droplet accumulation. Flow cytometry quantification (*left)* and representative histogram (*right*) of BODIPY+ cells in TSPO-KO and WT primary microglia stimulated with or without LPS. Dots represent individual wells (n = 9/group). (**B**) Representative confocal images of BODIPY (green) with DAPI (blue) staining in WT and TSPO-KO microglial cultures. (**C**) Flow cytometry quantification (*left)* and representative histogram (*right)* of BODIPY+ cells in WT primary microglia treated with TSPO ligand, Ro5-4864 (Ro5), or Vehicle. Dots represent individual wells (n = 9/group). (**D**-G) Flow cytometry quantification (**D, F**) and representative confocal images of Aβ uptake (**E, G**) in TSPO-KO and WT cultured primary microglia (**D, E**), and cultured primary WT microglia treated with TSPO ligand, Ro5-4864 (100nM), or Vehicle (1% Ethanol) (**F, G**). Dots represent individual wells (n = 9/group). *\*p* < 0.05, *\*\*\*\*p* < 0.0001. Abbreviations: MFI: Mean fluorescent intensity.

### TSPO modulates Aβ phagocytosis in cultured microglia

Next we investigated the effect of TSPO-KO on phagocytic functions in microglia, which play an important protective function in AD through clearance of Aβ. TSPO-KO microglia exhibited impaired phagocytosis of Aβ compared to WT microglia measured by flow cytometry (*t* test: CI= −45.3, −34.3; *p* = <0.001; n = 9/group, average of 3 experiments; Fig. 3D) and visually confirmed by ICC (Fig. 3E). Conversely, treatment with the TSPO ligand Ro5-4864 increased phagocytosis of Aβ in primary microglia measured by flow cytometry (*t* test: CI = 37.3, 2.42e+02; *p* = 0.0276; n = 9/group, average of 3 experiments; Fig. 3F), and confirmed by ICC (Fig.3G). We hypothesized that the phagocytic impairment in TSPO-KO microglia was driven by metabolic deficiency, with insufficient ATP production to support energy demanding functions such as actin polymerization required for phagocytosis. Consistent with this hypothesis, TSPO-KO microglia exhibited a marked reduction in actin levels (*t-*test: CI = −21.9, −9.39, p < 0.0001; n = 35 – 52 individual cells/group; Suppl. Fig. 1G). TSPO-KO did not affect cell viability in LPS or Aβ treated cultures (<1% propidium iodide+ cells in vehicle, LPS and Aβ treatment conditions measured via flow cytometry).

### TSPO is upregulated in phagocytic microglia, surrounding amyloid plaques in Alzheimer brain

To investigate if the functional association between TSPO and phagocytosis is consistent with the phenotype of TSPO expression in AD, we investigated TSPO expression in actively phagocytosing myeloid cells isolated from AD-mouse brain. Phagocytic cells were identified by the presence of internalized Aβ by flow cytometry (Fig. 4A - D). The proportion of Aβ^+^ phagocytes increased in APP-KI mice with age (One-way ANOVA; F _(2,16)_ = 126.931 *p <* 0.001; n = 4-7 mice/group; Fig 4A) and the majority were identified as microglia through expression of the microglia-specific marker TMEM199 (Fig. 4B). Aβ^+^ phagocytic microglia were characterized by elevated TSPO expression (Fig 4C), as well as upregulated expression of MHCII, CD68 and CD11c, markers previously associated with activated, phagocytic microglia phenotypes in aging and AD (Fig. 4C, D) (67). In contrast to Aβ^−^ microglia, Aβ^+^ microglia also exhibited high CD45 expression (Fig. 4D, E), consistent with previous findings that CD45^hi^ populations are the most phagocytically active mononuclear phagocytes in the brain (68). Expression of the microglial marker P2YR12 further corroborated the microglial origin of the Aβ^+^ phagocytes despite elevated CD45 expression. CD45, MHCII and CD11c expression progressively increased in Aβ^+^ phagocytic microglia as disease severity progressed in APP-KI mice between 5 and 10 months of age, perhaps reflecting the functional specialization of these microglia in response to Aβ pathogenesis (Fig. 4E).

**Fig. 4.**
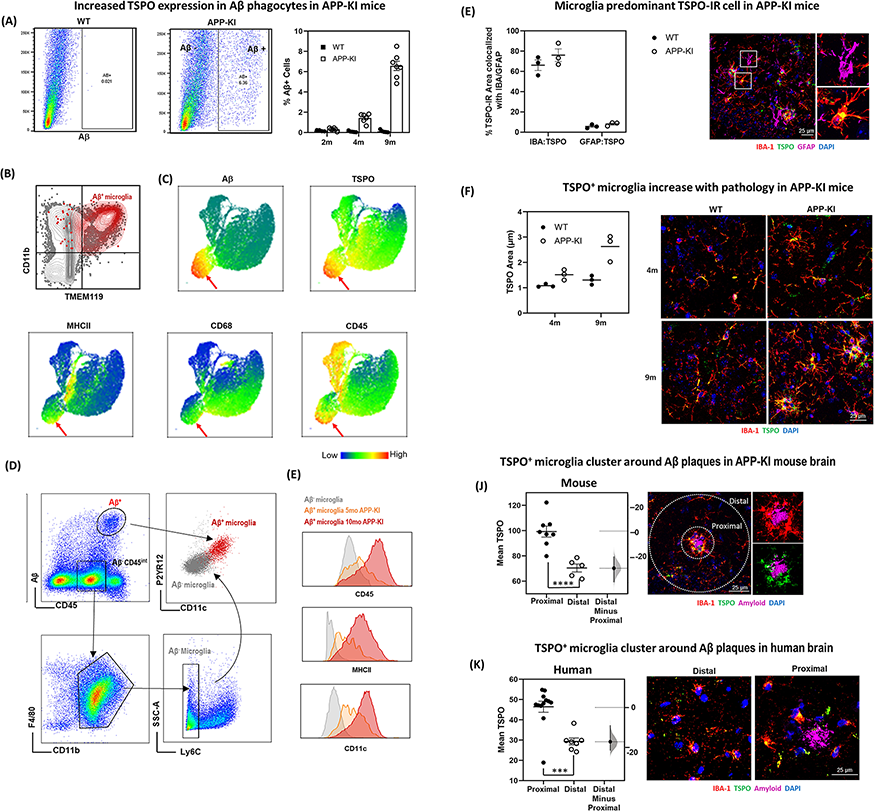
TSPO is upregulated in phagocytic Aβ^+^ microglia clustered around plaques in human and mouse AD brain. (**A**) Representative flow cytometry gating of Aβ+ phagocytes in WT (*left)* and APP-KI mice (*middle*). Quantification of Aβ+ phagocytes in APP-KI and WT mice at 2, 4, and 9m of age (*right)*. Dots in quantification graph represent individual mouse brains (n = 4-7/group). (**B**) Flow cytometry scatter plot showing CD11b and microglial marker, TMEM199 in Aβ^+^ cells, confirming the Aβ^+^ phagocytes are microglia. (**C**) Uniform manifold approximation and projection (UMAP) analysis showing 10,000 randomly sampled CD45^+^ cells from 12 month old APP-KI mice with distribution of Aβ, TSPO, MHCII, CD68 and CD45enriched populations indicated as a heatmap. Red arrow indicates Aβ^+^TSPO^hi^ population. (**D**) Representative flow cytometry histograms showing gating of CD45^hi^Aβ^+^ and CD45^int^Aβ^-^ microglial populations and relative expression of CD11c and microglial marker P2YR12. (**E**) Flow cytometry histograms showing CD45, MHCII and CD11c increase with age/disease progression in Aβ^+^ microglia (5 month vs 10 month old APP-KI mice). (**F**) TSPO colocalization with IBA-1^+^ microglia and GFAP^+^ astrocytes in WT and APP-KI mice at 9 months of age (n = 3/group). (**G**) Quantification of TSPO-IR in IBA-1^+^ microglial in WT and APP-KI mice at 4 and 9 months of age (n = 3/group). (**H**) Quantification and representative confocal images of TSPO immunoreactivity (TSPO-IR) in IBA-1+ microglia located proximal (10 μm) versus distal (80 μ m) to amyloid plaques in (**I**) APP-KI mice at 9 months of age (n = 5/group) and (**K**) human AD brain (n = 7/group). *\*\*\*p* < 0.001, *\*\*\*\*p* < 0.0001. Abbreviations: MFI: Mean fluorescent intensity.

Since previous studies have implicated microglia clustered at Aβ plaques in phagocytic clearance (69), TSPO expression was analysed by immunohistochemistry in mouse and human AD brain. In APP-KI mice, TSPO immunoreactivity was predominantly associated with microglial TSPO, with marked colocalization between TSPO and IBA1+ microglia in brain parenchyma, with some colocalization with GFAP-positive astrocytes (n = 3/group; Fig. 4F). Vascular and neuronal TSPO expression was also evident in both WT and APP-KI mice, although this did not change in a disease-dependent manner. Microglial TSPO immunoreactivity increased progressively between 4 and 9 months of age (Two way ANOVA; Main Effect (Age): F _(1,8)_ = 14.83 *p =* 0.049; n = 3/group; Fig. 4G) and TSPO expression was significantly higher in APP-KI mice compared to WT mice at 9m of age (Two way ANOVA; Main Effect (Genotype): F _(1,8)_ = 25.77 *p =* 0.001; Pairwise Comparison (APP-KI 9m v WT 9m): *p =* 0.0028). Significantly increased TSPO immunoreactivity was observed in IBA1^+^ microglia clustered around, and infiltrating Aβ plaques compared to microglia distal to plaques in both APP-KI mouse brain (*t* test: CI= −39.7, −19.8; *p* = 0.0002; n = 5/group; Fig. 4H) and human AD tissue (*t* test: CI= −21.6, −8.94; *p* = 0.0016; n = 7 AD brains; Fig. 4I).

### TSPO deletion impairs Aβ phagocytosis in Alzheimer’s mice and exacerbates pathology

Since our *in vitro* findings demonstrated TSPO regulates phagocytosis and we observed increased TSPO expression in Aβ^+^ microglial phagocytes in APP-KI mice, we examined whether TSPO knockdown affected the ability of these cells to phagocytose Aβ in APP-KI crossed with TSPO knockout mice (APP x TSPO-KO). APP x TSPO-KO mice exhibited a significant reduction in Aβ^+^ phagocytic microglia compared to age-matched APP-KI mice (*t* test: CI= −7.77, −3.71; *p* < 0.0001; n = 5/group; Fig. 5A), despite no effect of TSPO-KO on total microglial numbers measured by flow cytometry or IHC in APP-KI mice (all *p* > 0.911). Reduced Aβ^+^ phagocytic microglia numbers in APP x TSPO-KO mice was associated with increased amyloid load at 12 months of age, compared to APP-KI mice (Two way ANOVA; Main Effect (Genotype): F _(1,10)_ = 8.094 *p =* 0.0174; n = 3-4/group; Fig. 5B). No significant effect of TSPO-KO was observed on percentage of lipid-droplet high microglia in APP mice was also observed (*t* test: CI= −1.82, 8.43; *p* = 0.147; n = 4-5/group; Fig. 5C).

**Fig. 5.**
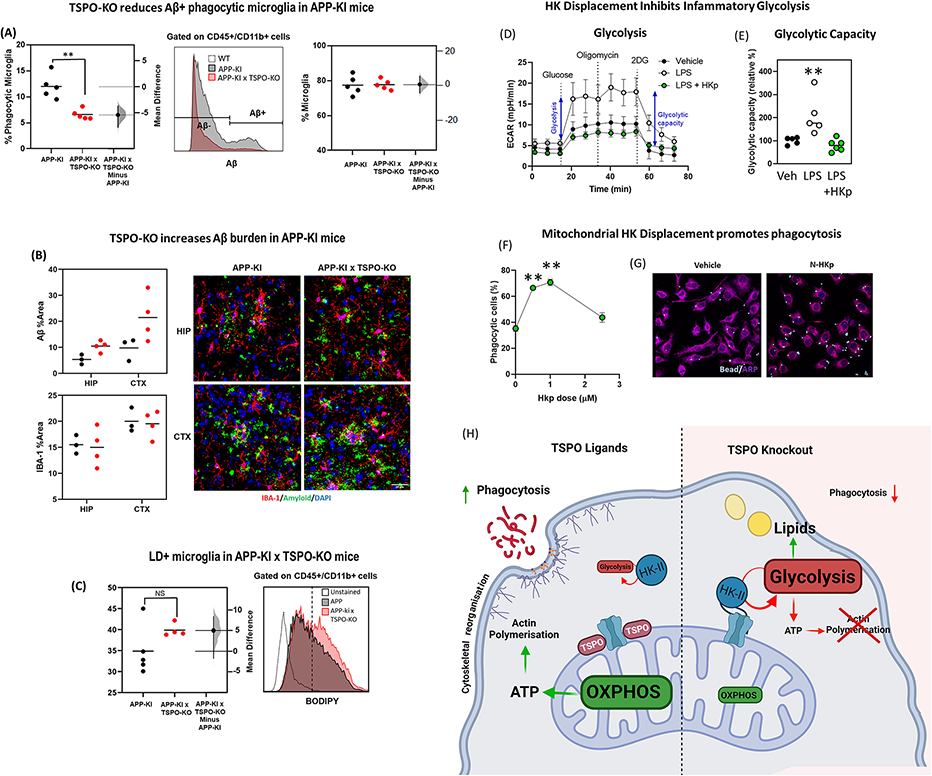
TSPO deletion impairs A β phagocytosis in Alzheimer’s mice and exacerbates pathology, mitochondrial hexokinase displacement promotes phagocytic function *in vitro*. **(A)** Flow cytometry quantification and representative histogram of Aβ microglial cells gated on CD45^+^/CD11b^+^ in APP-KI and APP-KI x TSPO-KO mice. Dots represent individual mouse brains (n = 4-5/group). (**B**) Quantification and representative confocal images of A β and IBA-1 immunoreactivity in APP-KI and APP-KI x TSPO-KO mice (n = 3-4/group). (**C**) Flow cytometry quantification and representative histogram of BODIPY+ microglia in APP-KI and APP-KI x TSPO-KO mice. (**D**) Glycolysis test in BV2 microglia treated with either vehicle or LPS +/- HKp showing mean ± SEM extracellular acidification rate (ECAR). n = 5-6 wells/group. **(E)** Quantification of glycolytic capacity calculated from glycolysis assay. Dots represent individual wells. (**F**) Phagocytic efficiency calculated as % cells ingested latex beads measured via flow cytometry in BV2 microglia treated with increasing doses of HKp (0 – 2.5 µM). Dots represent individual wells. n = 9/group, average of 2 independent experiments. (**G**) (**H**) Representative confocal images of phagocytic BV2 microglia treated with vehicle or HKp showing ingested beads (blue) and marker of cytoskeletal architecture (violet). (**I**) Schematic of proposed HK mediated control of microglial immunometabolic function. TSPO dampens mitochondrial HK recruitment, promoting OXPHOS, generating abundant ATP for energetic immune functions requiring cytoskeletal remodeling such as phagocytosis. TSPO-KO results in excessive mitochondrial HK recruitment, increasing glycolytic metabolism of glucose and storage of metabolized carbohydrates as lipids. Insufficient ATP production inhibits energetic immune functions such as phagocytosis. Created with BioRender.com.\**p* < 0.02, \*\**p* < 0.003, \**\*\*p* < 0.0001. Abbreviations: MFI: Mean fluorescent intensity.

### Mitochondrial hexokinase displacement reprogrammes microglial metabolism and promotes phagocytic function

We hypothesized that phagocytic impairment in TSPO-KO microglia was driven by metabolic deficiency resulting from excessive mitochondrial HK recruitment. Therefore we tested the effect of mitochondrial HK displacement on microglial glycolysis and phagocytosis. Mitochondrial HK was competitively displaced using a cell permeable, truncated N-terminal hexokinase peptide fragment lacking the enzymatic domain (HKp) (66). Hexokinase displacement with HKp reduced glycolytic capacity measured by extracellular acidification (ECAR) in LPS stimulated BV2 microglia, returning glycolysis to levels observed in unstimulated cells (one-way ANOVA: F _(2, 13)_ = 9.882, *p* = 0.0025; LPS vs LPS + HKp post-hoc: *p* = 0.003; n = 5-6; Fig 5D, E). HKp did significantly effect mitochondrial membrane potential measured by TRME (*t*-test: CI= 4.47e+03, 6.08e+04; *p* = 0.076; n = 6/group, average of two experiments; Suppl. Fig. 1H). Displacement of mitochondrial HK using N-HKp increased phagocytic efficiency dose-dependently in BV2 microglia (One-way ANOVA: F_(3,32)_ = 43.33, *p* < 0.0001; Control vs 0.5 µM or 1 µM post-hoc: *p <* 0.0001; n = 9/group, average of two experiments; Fig. 5F, G). These findings suggest mitochondrial HK may be targeted to regulate microglial immunometabolic function.

## Discussion

The mechanisms coordinating mitochondrial metabolism to fuel phagocytosis in AD remain poorly understood. Here we demonstrate that mitochondrial displacement of the glycolytic enzyme, hexokinase-II (HK) regulates microglial metabolism and phagocytosis, and that deletion of TSPO inhibits this. These findings provide new evidence that microglial immunometabolic programming is regulated via changes in subcellular localization of the key metabolic enzyme, HK, and that TSPO functions as an immunometabolic brake via regulation of mitochondrial HK recruitment, preventing hyperglycolysis and promoting phagocytosis in AD (Fig. 5H).

TSPO has been widely investigated as a biomarker of neuroinflammation in AD, and TSPO ligands exert protective effects by reducing Aβ accumulation in AD mouse models (10, 17). However, the precise immunophenotype associated with TSPO expression, and how TSPO exerts it’s anti-amyloidogenic effects are unknown. We provide novel evidence that TSPO plays a crucial role in promoting phagocytic clearance of Aβ via regulation of metabolic programming. In AD brains, TSPO-positive microglia clustered around Aβ plaques, and in AD mouse models phagocytic microglia were TSPO-enriched. These TSPO-enriched microglia were characterized by high expression of a number of markers previously associated with phagocytic microglial phenotypes including CD11c, MHII and CD68. These findings indicate that in AD, TSPO-PET signals are a marker of activated phagocytic microglia, and therefore may be useful for monitoring the efficacy of novel immunotherapeutics aimed at promoting phagocytic function. Supporting this, elevated TSPO-PET signals in prodromal AD have been associated with slower disease progression, suggesting a potential protective role of TSPO-associated inflammatory responses (70{10).

Elevated TSPO expression was observed in Aβ+ phagocytes isolated from APP-KI mice, which were identified as CD11c^+^ microglia. CD11c^+^ microglial subpopulations have been described across a number of studies in AD and ageing (also referred to as primed microglia, late response microglia, DAM, ARM, or MGnD; see (see 67 for full review), and these populations are characterized by enhanced capacity for uptake and lysosomal degradation of Aβ (71). CD11c combines with Integrin β2 (αxβ2; CD18) to form the complement receptor 4 (CR4), which is pivotal for phagocytosis (72), and, although CD11c is commonly used as a marker for peripheral dendritic cells, brain resident CD11c^+^ populations that express microglia specific markers TMEM199 and P2RY12 have also been identified (73). In line with this, Aβ^+^CD11c^+^ phagocytes isolated from APP-KI mice in the present study also expressed TMEM199 and P2RY12, confirming their status as brain resident microglia. Further, these cells exhibited increased expression of markers previously identified in phagocytic AD-associated microglial sub-populations, including MHCII, an antigen-presenting molecule involved in activating immune responses (74), and CD68, a lysosome associated marker involved in phagosome maturation (71).

Interestingly, TSPO deletion induced a microglial phenotype that overlapped with dysfunctional microglia observed in aging, characterized by metabolic changes, accumulation of lipid droplets and phagocytic impairment (65). Both lipid droplet accumulation and mitochondrial bioenergetic impairments have previously been identified as features of detrimental microglial sub-populations in aging and AD, characterized by impaired phagocytosis. For instance, Marschallinger et al. identified a unique sub-population of phagocytic-impaired microglia that exhibit increased lipid-droplet formation and increase with ageing and AD (65). Lipids can be synthesized from acyl-CoA generated from metabolism of carbohydrates such as glucose. In aging and disease this is linked to metabolic reprogramming from OXPHOS to glycolysis, which was recapitulated by TSPO deletion (63, 64). Therefore, we propose TSPO deletion induces a switch to conversion of glucose into lipids for storage, rather than mitochondrial energy production, resulting in insufficient energy supply to support energetically demanding processes such as actin polymerization for phagocytosis (see Fig. 5H).

Supporting this, in TSPO-KO mouse brain, transcriptional downregulation of genes involved in mitochondrial respiration and oxidative phosphorylation, as well as upregulation of genes associated with lipid synthesis were observed. This included *ACLY,* a key enzyme involved in the shift from carbohydrate metabolism via the TCA cycle to cholesterol and fatty acid synthesis. This is consistent with previous studies that have reported altered OXPHOS and glycolysis following stimulation with TSPO ligands (75) or TSPO deletion (20). Interestingly, TSPO has long been known to affect lipid biosynthesis and usage, specifically cholesterol, in other cells types, with a hypothesized role in steroidogenesis its most widely studied function. While recent genetic loss of function studies have shown that TSPO is not critical for steroidogenesis in the periphery (76, 77), TSPO deletion drastically reduced steroidogenesis in the brain (78). Although microglia are not steroidogenic cells, our findings suggest TSPO may play a role in regulation of the synthesis of lipids including cholesterol from carbohydrates. Therefore our findings may offer a potential alternative explanation of the effect of TSPO on steroidogenesis via effects on lipid synthesis from carbohydrate substrates to alter cholesterol bioavailability for steroidogenesis. This hypothesis could be tested in future studies.

Proteomic analysis of the TSPO interactome network identified the 14-3-3 family scaffold adaptor and chaperone proteins, as a hub in the TSPO-interactome network. The 14-3-3 adaptor proteins are metabolic regulators (59) and have been previously shown to interact with the TSPO-complex in Leydig cells, with 14-3-3 adaptor phosphoserine protein binding motifs identified on functionally important sites on both TSPO and VDAC (60, 61). Our findings indicate these adaptor proteins may also be key components of the functional TSPO complex in the brain. Another previously reported TSPO interactor identified in the brain TSPO interactome included ATP synthase, which has also been previously shown to be regulated by the 14-3-3 adaptor proteins. TSPO interactions with the 14-3-3 adaptor proteins and ATP synthase may potentially explain how TSPO promotes OXPHOS. In contrast, we demonstrate that TSPO deletion promotes mitochondrial recruitment of the glucose metabolising enzyme, HK. In other cell types mitochondrial HK recruitment drives glycolytic metabolism. We propose that TSPO therefore coordinates the balance of OXPHOS and non-oxidative glucose metabolism in microglia. Importantly, these findings indicate that targeting mitochondrial HK may offer a novel immunotherapeutic approach to promote microglial phagocytosis in AD.

## Acknowledgements

The authors are grateful to Drs Makoto Higuchi and Bin Ji (National Institutes for Quantum and Radiological Science and Technology, Japan) for providing the TSPO-KO mouse model, and Drs Takaomi Saido (RIKEN Center for Brain Science) and Takashi Saito (Nagoya City University Graduate School of Medical Sciences) for providing the APP-KI mouse model, and Jianpeng Sheng (SBS, NTU) for technical assistance. This project was funded by the Nanyang Assistant Professorship Award from Nanyang Technological University Singapore (AMB).

## Author contributions

Conceptualization: AMB; Study design: LHF, KOL, AMB; Methodology: All Authors; Formal analysis: LHF, KOL, GDA, SL & AMB; Writing: LHF, KOL & AMB; Visualization: LHF, KOL & AMB; Supervision: AMB; Funding acquisition: AMB. All authors approve the manuscript for publication.

---

**Supplementary Fig. 1.** (**A**) Mitochondrial membrane potential (MMP) measured by TRME fluorescence in WT and TSPO-KO microglial cultures. n = 4 – 6 wells/group. (**B, C**) Quantification of mitochondrial footprint (**B**) and fragmentation (**C**) in WT and TSPO-KO microglia. Dots represent individual cells (n = 40/group). (**D**) Representative confocal images of mitochondria (ATPB, red) in single WT and TSPO-KO microglial cells. (**E**) Flow cytometry gating of BODIPY^+^ CD11b^+^ cells in LPS treated WT mice. Quantification of TSPO levels in lipid droplet^+^ (LD^+^) and LD^-^ microglia (CD11b^+^ cells). n = 3 mice. (**F**) Flow cytometry quantification and representative histogram of PLIN2 expression in phagocytic (+) and non-phagocytic (-) microglia. Abbreviations: AU: Arbitrary units; SSC-A: side scatter; FSC-A: forward scatter. (**G**) Quantification of F-actin content measured using phalloidin. Dots represent individual cells. n = 35 - 52/group. Representative confocal images of phalloidin stained F-actin in primary WT of TSPO-KO cultured microglia. LUT (mpl-inferno) of confocal images represents F-actin intensity. (**H**) MMP measured by TRME fluorescence in BV2 microglial cultures treated with vehicle (veh) or HKp. FCCP shown as positive control for MMP depolarization. n = 4 – 6 wells/group. \**p* < 0.05.

**Supplementary Table 1. Details of antibodies used for flow cytometry experiments.**

**Supplementary Table 2. Demographic data and neuropathological features of AD cases.** Abbreviations: AD Alzheimer’s disease, F female, M male, CAA cerebrovascular angiopathy. CERAD scores (79) for neuritic plaque densities: 0 = none (no plaques), 1=sparse, 2=moderate, and 3=High density.

**Supplementary Table 3. Significant differentially expressed genes in hippocampus of WT versus TSPO-KO mice in control and LPS stimulated conditions.** LFC > 2, FDR < 0.05.

**Supplementary Table 4. Functional Gene Set Enrichment (FGSEA) comparing transcriptional responses in LPS-stimulated TSPO-KO and WT mice.** Top 5 significantly functionally enriched pathways in the following categories shown: (1) immune and inflammatory pathways, (2) lipid metabolism, and (3) mitochondrial metabolism.

**Supplementary Table 5. Candidate TSPO interactors.** Significantly enriched proteins identified by IP-MS in WT and APP-KI brain relative to TSPO-KO background. Fold Change > 2; FDR < 0.1.

**Supplementary Table 6. TSPO interactome network functionally enriched pathways (KEGG & Reactome).** FDR < 0.05.

