## Supplementary figures and images for "Mitochondrial control of microglial phagocytosis in Alzheimer’s disease"

### Supplementary Figure 1

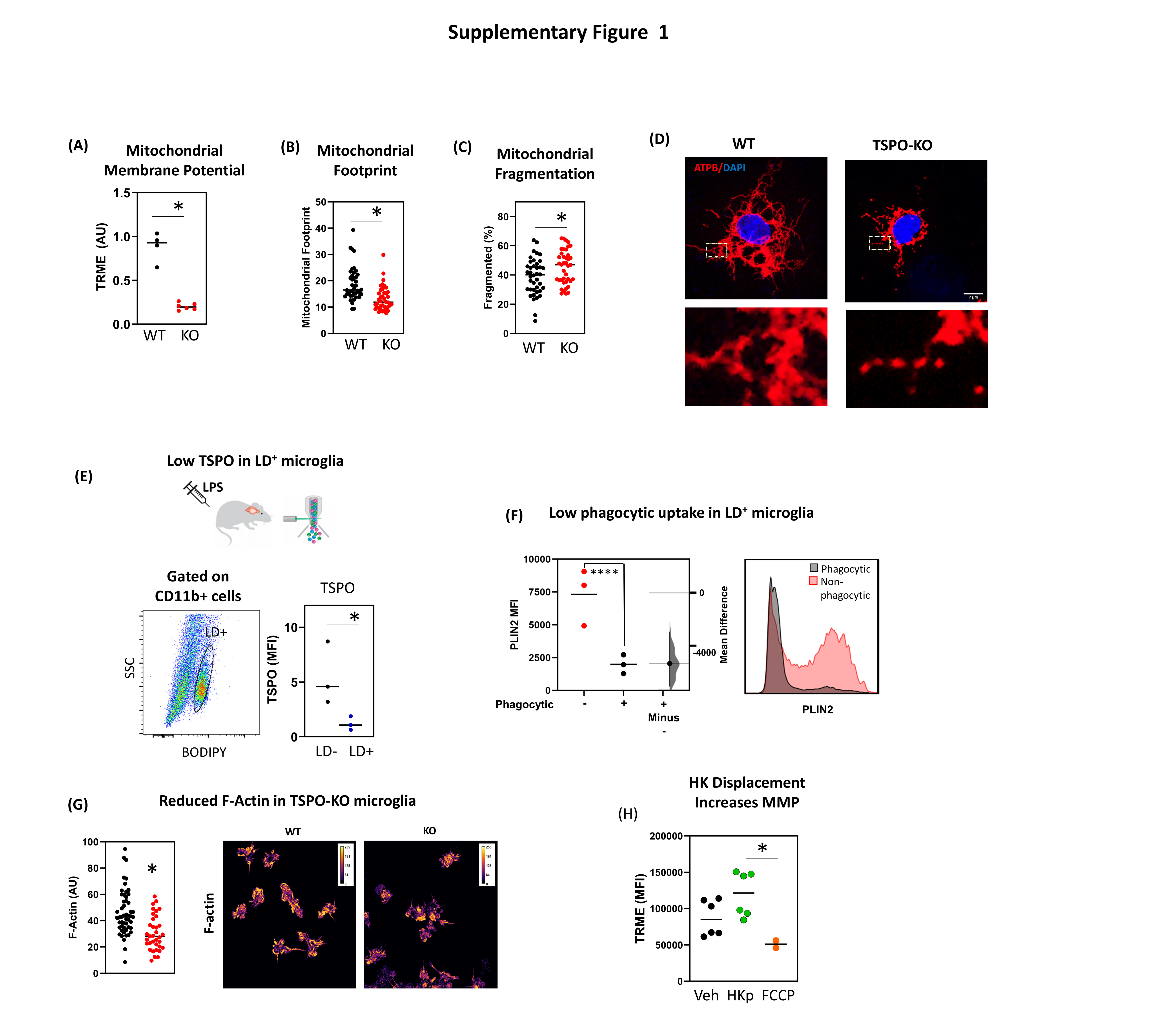
