## Supplementary Table 1 for "Mitochondrial control of microglial phagocytosis in Alzheimer’s disease"

Supplementary Table 1: Cohort characteristics

| Case number | Age  (years) | Sex | Braak stage^[1]^ | CERAD score^[2]^ | Clinical diagnosis | Post-mortem delay (h) |
| --- | --- | --- | --- | --- | --- | --- |
| 1  2  3  4  5  6  7 | 68  84  80  87  62  90  89 | M  F  F  M  M  M  F | 4  4  4  6  6  3  4 | 2  3  3  3  3  2  2 | AD  AD, Dementia  AD, CAA  AD  AD, CAA  AD  AD | 24  13.5  28  23  21  30.5  26.5 |

*AD* Alzheimer’s disease, *F* female, *M* male, *CAA* cerebrovascular angiopathy. CERAD scores for neuritic plaque densities: 0 = none (no plaques), 1 = sparse, 2 = moderate, and 3 = High density.
