## Supplementary methods for "Mitochondrial control of microglial phagocytosis in Alzheimer’s disease"

**Human post-mortem tissue samples**

The Multiple Sclerosis and Parkinson’s Tissue Bank at Imperial College London provided all AD post-mortem tissues for this study. Fully informed consent was obtained for the post-mortem donation under ethical approval by the National Research Ethics Committee (08/MRE09/31) and use of the tissue was approved by the NTU Institutional Review Board (IRB-2018-09-052-03). Neuropathological analysis confirmed AD pathology. The demographic data and neuropathological features of the AD cases are shown in Suppl. Table 1. For immunofluorescence analysis, 10 µm thick snap-frozen sections from 7 AD cases (4 males, 3 females; median age of death = 84 years, range = 62‒90 years; Braak stages III‒VI (Braak and Braak, 1991); median port-mortem delay = 24 h; range = 12‒28 h) were cut from the hippocampus/entorhinal cortex region based on tissue availability.

**RNAseq data generation & analysis**

Frozen hippocampi were dissected out of hemibrains at -20˚C, homogenised immediately in Trizol and total RNA extracted. Briefly, an aqueous phase was induced with the addition of 1:5 volume of chloroform to the Trizol and centrifugation for 12000 rpm, 15 min. The aqueous phase was then extracted and transferred into a Qiagen RNeasy mini kit column and the total RNA was extracted according to manufacturer’s instruction. Quality and purity of the total RNA was checked using the Agilent 2100 Bioanalyzer with Agilent 6000 Nano Kit. Total RNA of the samples had a minimum of RIN 8.3 with an average of RIN 9 and above.

**Immunoprecipitation mass-spectrometry (IP-MS)**

Frozen mouse hemibrains were homogenised within HEPES 5% sucrose buffer then solubilised with 0.1% Triton X-100 for 1.5 hrs and the soluble fraction were extracted and precleared with IgG Rabbit isotype control (Abcam ab172730) and Protein A/G agarose (Santa Cruz sc-2003) beads for 4 hrs. Soluble precleared fractions were then incubated overnight with rabbit anti-TSPO antibodies (Abcam 109497). Samples were then further incubated with 30 uL of Protein A/G agarose beads (Santa Cruz sc-2003) and subsequently washed three times with HEPES sucrose buffer. Immunoprecipitated proteins were eluted by incubating the beads with 2 $\times$ SDS laemlli buffer for an hour at 37°C. Eluted proteins were run on a 5% stacking 12% resolving gel on constant voltage 70V. Gels were fixed in 50:40:10 methanol: acetic acid: water for 30 min before colloidal blue staining. Gels were destained and gel fractions with stained bands were excised and prepared for LC-MS.

LC-MS and peptide detection was provided by Proteomics and Mass Spectrometry Facility in School of Biological Sciences, Nanyang Technological University. Briefly, the peptides were separated and analyzed using a Dionex Ultimate 3000 RSLCnano system coupled to a Q Exactive instrument (Thermo Fisher Scientific, MA, USA). Separation was performed on a Dionex EASY-Spray 75 μm × 10 cm column packed with PepMap C18 3 μm, 100 Å (Thermo Fisher Scientific) using solvent A (0.1% formic acid) and solvent B (0.1% formic acid in 100% ACN) at flow rate of 300 nL/min with a 60 min gradient. Peptides were then analyzed on a Q Exactive apparatus with an EASY nanospray source (Thermo Fisher Scientific) at an electrospray potential of 1.5 kV. Raw data files were processed and searched using Proteome Discoverer 1.4 (Thermo Fisher Scientific). The Mascot algorithm was then used for data searching to identify proteins.

**Cell Culture and treatments**

The immortalized murine microglial BV-2 cell line was grown and maintained in complete Dulbecco’s modified Eagle serum (DMEM) containing 10% fetal bovine serum (FBS) and 1% penicillin-streptomycin (DMEM-COM) at 37°C in 5% CO_2_. The BV-2 cell line was used because they have been shown to recapitulate inflammatory responses of primary microglia (Henn et al., 2009), and demonstrate a robust phagocytic response (Koenigsknecht and Landreth, 2004, Mandrekar et al., 2009).

Primary microglia were derived from the brains of neonatal mice as previously described (Lian et al., 2016). Briefly, postnatal day 1-3 mice were decapitated, their brains were dissected and meninges and blood vessels were removed completely. Tissues were transferred to 50 mL tubes and dissociated in 1.5 mL 2.5% trypsin at 37°C for 10 min. Dissociation was terminated by adding an equal volume of DMEM-COM, followed by 750 μL 10 mg/mL DNase. Tubes were then centrifuged at 400$\times$g for 5 min, the supernatant was removed and pellet was triturated in warm DMEM using a 1 mL pipet tip. Cells were plated in 75 mm flasks coated with poly-L-lysine (0.1 mg/mL; Sigma Aldrich) at a density of 5 × 10^7^ cells per flask. Media was replaced the next day with DMEM-COM. Cells were grown for 5 d without changing the medium to allow microglial proliferation. Microglia were harvested by vigorously tapping the flasks for 5 min. Purity of cultures was determined by staining with the microglial markers CD11b and CD45, and analysed by flow cytometry. All primary cultures were determined to be > 99.9% pure microglia.

**Mitochondrial bioenergetics in primary microglia**

Mitochondrial bioenergetics in cultured microglia were measured using the XF Cell Mito Stress Test Kit (#103015-100, Seahorse Biosciences, North Billerica, MA) according to methods described in the XFe96 Extracellular Flux Analyzer User Manual. Primary microglia were plated into XFe96 cell culture plates at a density of 50,000/well in 200 μL of DMEM-COM (with 0.5 ng/mL GM-CSF) and allowed to adhere overnight in 37 °C incubator with 5% CO_2_. XF sensor cartridges were hydrated with sterile water and kept at 37°C overnight in non-CO_2_ incubator. Prior to experiments, the sensor cartridges were hydrated with 200 μL XF Calibrant and kept at 37°C in non-CO_2_ incubator for 45 – 60 min and the injection ports were then loaded with test compounds. For Mito Stress experiments, following cell adherence, the media was removed and replaced with assay media XF RPMI medium (pH 7.4) (#103576-100) supplemented with 1 mM pyruvate (#103578-100), 2 mM glutamine (#103579-100), and 10 mM glucose (#103577-100). The cells were incubated at 37 °C in non-CO_2_ incubator for 60 min prior to assay. OCR was measured by sequential injection of oligomycin (1.5 μM final concentration), FCCP (2 μM final concentration) and Rotenone/Antimycin A (0.5 μM final concentration).

**Isolation of myeloid cells from mouse brain**

Isolation of myeloid cell populations was performed as previously described (Sheng et al., 2015). Briefly, hemibrains from adult APP-KI, APP-KIxTSPO-KO, and age matched WT controls were minced and digested in IMDM (2% FBS) containing 100 mg/mL Collagenase (Roche Applied Science, Basel Switzerland), 1.2 U/mL Dispase (Roche Applied Science, Basel Switzerland), and 20 Units/mL DNase I (Life Technologies) for 30 min at 37°C on a shaker. Tissue suspensions were passed through a 19-guage syringe and filtered through a 40 μm cell strainer to obtain homogenous suspensions. Cell suspensions were centrifuged at 350g for 5min at RT, the supernatant discarded, and cell pellets resuspended in 4 mL 40% Stock Isotonic Percoll (SIP) (90% Percoll + 10% 10x phosphate buffered saline or PBS (GE Healthcare, Life Sciences) per brain and overlaid on 3 mL 70% SIP and then centrifuged for 10 min at 2800 rpm, with deceleration 4 at RT. Myelin was aspirated and the cell solution was collected from the 70%-40% interphase layer, centrifuged at 600g for 5 min and the pellet was resuspended in 2 mL PBS (2% FBS), ready for staining with fluorescence cytometry antibodies.

**Image collection and analysis in *in vitro* studies**

For analysis of mitochondrial footprint, fragmentation and HK binding, images were captured at 63x magnification and z-stacks were obtained based on optimised Nyquist sampling (0.21 µm ~0.33 µm step size) using a confocal microscope (Zeiss Laser Scanning Microscope-800 upright microscope) with Zeiss ZEN imaging software. Image analysis was performed in a user-blinded manner using Imaris 9.2.0 Image Visualization and Analysis Software. Specifically, mitochondria surfaces were background subtracted and created with smooth detail of 0.125μm. Mitochondria were classified as fragmented (>0.8 sphericity), tubular (<0.8 sphericity and volume <5μm) and elongated mitochondria (<0.8 sphericity and vol >5μm). Representative figures of mitochondria staining (ATPB and HK) were processed as previously described (Chaudhry et al., 2020 doi: <https://doi.org/10.1152/ajpendo.00457.2019>) and are presented as Maximum Intensity Z- Projections.  Respective raw images (xy drift corrected) were used for all intensity quantifications. Same pre-processing steps have been performed for each representative image comparison set. Using ImageJ/Fiji, representative image of mitochondrial ATP has been processed using the following commands: 1) “Single Substack” or Slice of highest mitotracker intensity was extracted from the z-stack of each image. 2) “Enhance Contrast (saturation=0.35)” to increase contrast of image. 3) “Unsharp mask radius =1 mask=0.60” to sharpen image visualisation. 4) “Median.., radius=1” to remove salt and pepper noise of the image. Kymograph was created using line tool from Fiji and straighten option. Representative image of Hexokinase colocalization to mitochondria has been processed using the same steps as Mitochondrial ATP excluding the first step. Background was subtracted using the “rectangle roi” values and rolling ball =10 pixels. Colocalization diagram was generated using Fiji Plugin Fret and Colocalization Analyzer.
